# AI for IACUC: Accurate Initial Assessment of Institutional Animal Care and Use Committee Protocols

**DOI:** 10.1101/2025.08.22.671834

**Authors:** Brent Vasquez, Jeremy DeRicco, Bill J. Yates, Robert K. Cunningham

## Abstract

Every animal protocol requires careful review to ensure humane principles, laws, and policies are followed before any experiments may be performed. Developing protocols and receiving approval can be a lengthy process requiring multiple rounds of feedback and refinement. Artificial Intelligence (AI) has the potential to improve the quality and speed of IACUC protocol reviews by detecting submission errors. We describe an approach to achieve this and show that our approach can be replicated, while addressing issues of ethics and bias, robustness, and trustworthiness^1^. Replication is addressed by developing a template and demonstrating that it can be reused. Ethics and bias issues are addressed by early, limited use of a large language Model (LLM) to identify errors and rapidly provide useful feedback to the protocol developer. In our approach, subsequent steps still require a review by the IACUC board, which retains ethical responsibility for approval. Robustness and trustworthiness were evaluated by using the template that is replicated for eleven distinct checks and performed on 50 new protocols using two different LLMs. We show that every actual problem in those submissions (within the test parameters) is found by our AI review (i.e., recall=100%), giving IACUC boards confidence in the utility of our system. In addition, precision for our checks is between 80%-100%, with most being 100%, thereby making the system a practical compliment to administrative protocol review for those submitting. Overall, we report F1 scores of between 89-100%.

## Introduction

Investigators performing animal and human research are obligated to conduct their studies ethically and follow regulatory requirements. Russell and Burch described the widely followed “3R principles” of performing humane experimental studies on animals – *Replacement:* seek alternatives to using animals in research; *Reduction:* minimize the number of animals needed; and *Refinement:* modify experimental procedures to reduce pain and distress experienced ^2^. In the United States, the Animal Welfare Act of 1966 ^3^ and the US Public Health Service Policy ^4^ require an Institutional Animal Care and Use Committee (IACUC) to prospectively evaluate study protocols to ensure compliance with ethical and regulatory requirements. In the US, institutional review boards (IRB) perform a similar function for human subjects^5^. Many other countries have similar regulations ^6,7^.

Large numbers of experiments on animals are carried out each year. For example, in the European Union, more than 4 million procedures were performed on mammals in 2018^8^, each requiring careful review and assessment. Tools and processes that speed review and assessment while ensuring humane treatment are therefore valuable.

In the United States, IACUC reviewers examine carefully designed and described protocols to ensure compliance with applicable law and policies. Each protocol includes a justification of the species and number of animals used, specific experimental procedures employed, and a pain management and monitoring plan^9^. It is common for junior researchers to develop research protocols in collaboration with a more senior researcher, with the junior member documenting the procedures in forms. Inconsistencies and omissions within the IACUC protocol delay approval and thus the start of experiments. Such errors can also contribute to an increased workload for the IACUC, which is often required to review a high volume of protocols. For instance, between July 2023 and June 2024, the University of Pittsburgh’s IACUC conducted approximately 500 de novo protocol reviews, including both new submissions and three-year *de novo* reviews. Given the substantial review burden, implementing a systematic process automatically detecting inconsistencies and errors within protocols would be highly beneficial.

The protocol review process consists of multiple stages and typically requires several weeks. Below, we outline the process implemented at the University of Pittsburgh (Pitt), though similar procedures are followed at other institutions. Before initiating an experiment, the principal investigator (PI) submits a draft protocol via the Animal Research Online (ARO) system, Pitt’s implementation of the IACUC component of the Huron Research Suite (Huron Consulting Group, Chicago, IL). Pitt’s Animal Research Protections (ARP) Division supports the IACUC by conducting preliminary reviews of submitted protocols and providing feedback to PIs for necessary revisions. Once the PI addresses the initial comments and refines the draft, ARP facilitates the designated member review (DMR) process by coordinating communications between the PI and the IACUC. In cases where protocols contain errors or omissions, multiple rounds of review may be required to ensure accuracy and compliance with ethical and regulatory standards before final approval is granted.

The recent emergence of large language models (LLM) ^10,11^, and their demonstrated ability to both read natural language explanations and write accurate answers to well-documented questions^10^ suggests that they may be helpful for components of the review process^12^. However, these systems are not perfect. They have been shown to create confabulations and fabrications across a wide range of fields^13^. An open question is whether it is possible to develop a robust, reliable, ethical administrative tool for the animal protocol review process.

There are three common issues in identifying good tests for the use of LLMs – ethics and bias, robustness, and trustworthiness^1^. In this paper we address ethics and bias by carefully selecting how we use an LLM in the review process, using the LLM to review the initial submission and quickly identify a limited set of problems and to communicate that information efficiently to the researcher. We identified only initial administrative information to be reviewed by the AI, with final review and approval performed by the required legally constituted IACUC. We address robustness by evaluating our approach on two different large language models. For this study we used Gemini 1.5 Pro^14^ and a university-hosted version of ChatGPT-4o^15^, each of which allows the university to control data distribution and prevent accidental leakage of newly developed protocols. We address trustworthiness by gathering a large number of animal procedures and demonstrate excellent precision/recall/F1 performance across all procedures on both systems.

## Results

LLMs were programmed using prompt engineering on eleven IACUC protocol samples collected from July to August 2024. After extensive experimentation, it was determined that LLMs could be programmed to give reliable results using a prompt pattern^16^ template. Elements of the pattern for each prompt include: *output customization* – the LLM is instructed to take on a *persona* - “reply in a polite and professional tone,” and to provide a *visualization generator* by “marking each correct prompt with a green check mark and each incorrect prompt with a red ‘X’”. We then tasked the LLM to “Tally up all correct (green check marks) and incorrect responses (red ‘X’s).” The next element is *context control –* the LLM is provided the context of the query, by referencing the relevant paragraph or paragraphs from The Guide for the Care and Use of Laboratory Animals^4^. And finally, we include *the question itself*, often written in the style of a standardized test question. We found that the exact formatting of the question affected the performance of the system. In particular, it was important to separate each section with a paragraph break. A single example is provided in Table 1; material between paragraph breaks 1 and 4 are repeated for each additional prompt. All prompts are provided as supplementary material.

**Table 1:** Prompt Pattern Example.

| Structure Details | List structure |
| --- | --- |
| *Output customization:*<br>*persona* | Write in a polite and professional manner. |
| [Paragraph Break 1] |  |
| Question context control | Public Health Service (PHS) funding sources include: Administration for Children and Families (ACF); Administration on Aging (AoA); Agency for Healthcare Research and Quality (AHRQ); Agency for Toxic Substances and Disease Registry (ATSDR); Centers for Disease Control and Prevention (CDC); Centers for Medicare & Medicaid Services (CMS); Federal Occupational Health (FOH); Food and Drug Administration (FDA); Health Resources and Services Administration (HRSA); Children's Hospital (CHP foundation); Indian health Service (HIS); National Institutes of Health (NIH); Substance Abuse and Mental Health Services Administration (SAMHSA); and the Public Health Service Commissioned Corps. |
| [Paragraph Break 2] |  |
| Question Details and Instructions | If the only funding source checked in the document is a PHS funding source, then there should be a 'yes' response under "Is this study funded in part or whole by a PHS Agency?". If both a PHS funding source and "Internal" are checked, then the response under "Is this study funded in part or whole by a PHS Agency?" should be 'yes'. If the funding source mentioned in the funding source title table under section 1.3 is not a PHS agency, then there should be a 'no' response under "Is this study funded in part or whole by a PHS agency?" |
| [Paragraph Break 3] |  |
| Output customization | If this is correct put a green check mark, if this is incorrect put a red 'x'. |
| [Paragraph Break 4] |  |
| Output customization:<br>visualization generator | Tally all correct and incorrect prompts. |

Eleven different prompts were developed to address common errors that Pitt’s IACUC coordinators reported during the initial review process. Some prompts can be categorized as identifying errors completing the protocol form. We explored three types:

1. *Missing information* – the LLM is asked to determine if a response is missing in a specific section of the protocol,
2. *Impermissible information* – the LLM is asked to determine if something that may not be included (e.g., cost) was included, and
3. *Mismatched sections* – the LLM compares multiple related sections for inconsistencies.

Some of these prompts are simple such as assuring that an investigator listed in one section had a role specified in another. Other prompts are more complex and require computations and comparisons. For example, one prompt tasked the LLM to verify that the total number of animals requested for all experiments does not exceed the total number of animals requested for the study.

Other prompts required regulatory information about animal research and identified cases where the protocol failed to follow *legal requirements, ethical guidelines*, or *local policy*. Examples include appropriate species use, proper use of pain medication, and proper diet throughout the experiment.

After all queries achieved 100% precision and recall on all training data, 50 additional procedures were collected for testing. We tested on two different LLMs – PittGPT, which uses OpenAI’s ChatGPT-4o as its LLM, and Google’s Gemini 1.5 Pro. Performance was excellent - recall for each prompt was 100%, and the precision and F1 scores are depicted in Table 2. Prompt numbers in Table 2 reference those included in the supplementary material. The performance of the prompts was highly robust, with PittGPT achieving a recall of 1 and a precision of 0.98, while Gemini 1.5 Pro demonstrated a recall of 1 and a precision of 0.97. Despite being developed independently by two different companies, OpenAI and Google respectively, both models exhibited comparable levels of accuracy and effectiveness.

**Table 2.** Precision and F1 Scores for Each Type of Prompt Across All Protocols.

| Type | Prompt | Precision |  | F1 Score |  |
| --- | --- | --- | --- | --- | --- |
|  |  | PittGPT | Gemini 1.5 Pro | PittGPT | Gemini 1.5 Pro |
| Missing Information | 1 | 100% | 100% | 100% | 100% |
|  | 5 | 100% | 100% | 100% | 100% |
| Impermissible Information | 4 | 100% | 100% | 100% | 100% |
| Mismatched Sections | 2 | 100% | 100% | 100% | 100% |
|  | 3 | 100% | 98% | 100% | 99% |
|  | 7 | 100% | 100% | 100% | 100% |
| Identification Of Cases That Fail to Follow Standards | 6 | 80% | 80% | 89% | 89% |
|  | 8 | 93% | 84% | 96% | 91% |
|  | 9 | 100% | 100% | 100% | 100% |
|  | 10 | 100% | 98% | 100% | 99% |
|  | 11 | 100% | 100% | 100% | 100% |
| **Aggregate** |  | **98%** | **97%** | **99%** | **98%** |

The output generated by both systems was structured to provide clear and intuitive understanding of corrections needed in the protocol. Prompts that affirmed correct responses were marked with green check marks, while those that detected issues were provided with red “X” symbols. Additionally, the output from both LLMs included a concise explanation of each response, detailing the reasoning behind its classification as correct or incorrect. These explanations were presented professionally and courteously to ensure clarity and ease of interpretation. Results were obtained quickly; on average, responses take 8 seconds to process all 11 queries.

## Discussion

There are three common issues in identifying good tests for using LLMs – ethics and bias, robustness, and trustworthiness.

To address ethics and bias, we limit the use of LLMs to the first stage of evaluation (pre-screening of protocols), to provide rapid feedback, thereby reducing human workload during the most tedious part of assessment. The protocol is reviewed by experts only after all common errors have been identified and rectified. At this stage, the experts apply their knowledge to detect complex regulatory, research, clinical, and ethical issues that may not have been addressed through the initial error-detection process.

To address trustworthiness, we assessed precision, recall and F1 scores for our queries on a large sample of protocols. To test robustness, we measured the performance of our queries on two independently developed LLMs. We showed that when properly programmed, LLMs can accurately detect errors in animal use protocols. We developed a formula and structure to write new questions effectively. Simple prompts, like matching names in different sections or verifying adherence to euthanasia and diet guidelines, achieved 100% precision-recall scores demonstrating the LLMs proficiency in basic information extraction and comparison within the protocols. Prompts that compared animal numbers, verified PHS funding sources, and validated genotyping guidelines, also achieved near-perfect scores showcasing the LLMs ability to gather information across different protocol sections. Lastly, prompts for interpreting information within tables were the most challenging for both LLMs. Specifically, the outcome of prompts such as proper identification of pain and distress classifications, and classification of drugs and chemicals achieved precision scores at 80% and above. We attribute this to the table representation – in our examples, position alone indicated that a value was in a specific cell, rather than lines demarking cell boundaries. The varying complexities of the types of prompts clarify that both PittGPT and Gemini 1.5 Pro can execute demanding tasks.

Of note, newer versions of Gemini were released in 2025, but all tests were performed using Gemini 1.5 Pro. LLMs are expected to increase in performance over the years exponentially ^17^.

The success of the LLM performance suggests that both PittGPT and Gemini 1.5 Pro provide whole system opportunities. LLMs can be applied to other internal review processes by adapting the structure of prompts. In particular, PittGPT and Gemini 1.5 Pro can be useful for determining common factual errors in reports and can automate compliance checks by identifying deviations from regulations or internal policies, ensuring consistency and reducing human error^18^. We can also apply our methodology for ARRIVE (Animals in Research: Reporting In Vivo Experiments) to follow recommendations for proper vertebrate animal research. To fully ensure that the ARRIVE guidelines are properly addressed the methodology of our design of a prompt template can be used, but the technique needs to be adapted. By automating administrative checks, LLMs allow human reviewers to iterate on their checklists and fix problems in real time^19^. Additionally, they can personalize feedback for applicants, offering constructive criticism and improving the overall applicant experience. LLMs can help researchers by automating the process of analyzing large volumes of textual data to identify recurring themes, patterns, and trends^20^. This helps save significant time and effort, enabling deeper insights into complex datasets.

Finally, some LLM system agreements permit the provider to collect and save input for use in future training. Since protocols may contain sensitive or proprietary information, one must review service agreements carefully to ensure such information is not retained or misused, which should limit the use of some systems in conducting protocol reviews^21^. While the version of Gemini Pro used in this study had appropriate protections, OpenAI’s version of ChatGPT did not. This is one reason that Pitt developed PittGPT.

In summary, although most aspects of IACUC protocol review require careful thought, committees composed of members with multiple areas of expertise, and thorough IACUC deliberation, LLMs can be strategically incorporated to streamline review processes by accurately identifying common factual errors. Carefully engineered prompt patterns can ensure reliable AI-generated responses and can be adapted for various applications. The effectiveness of our approach highlights broader system-wide opportunities for AI integration and encourages further research into their evolving capabilities.

## Methods

### Formulation of Queries

Protocol review coordinators from the University of Pittsburgh Animal Research Protections (ARP) identified problems that are regularly encountered during the initial review of protocols. Problems were assessed in aggregate to identify possible approaches, and initial queries were drafted. Eleven previously approved protocols, sampled from submissions made from July to August 2024, were obtained from the University of Pittsburgh’s IACUC protocol processing system, Animal Research Online (ARO). ARO is a component of a legacy version of the Huron Research Suite (Huron Consulting Solutions). Protocols were exported into PDF format, which was subsequently ingested by the LLM systems.

### Defining and Refining Queries

Test protocols including intentionally introduced errors were created from eleven IACUC protocol PDFs for testing each newly created question, resulting in 11 new and 22 total examples. Each PDF was derived from one of the eleven exported PDFs and the protocol information related to the test queries were modified using Adobe Acrobat Pro (2024) version 24.003. Examples of modifications include redaction/addition of staff, materials, chemicals, drugs, and diet, modifications to animal sample size, and change in placement of check marks within tables.

Query development proceeded by uploading each PDF into one of two LLMs, each derived from the modern Transformer models^10^. We first selected Google’s Gemini 1.5 Pro-002^22,23^, a multimodal (i.e., text/audio/image/video in and text/audio/image out) model that demonstrated excellent performance across a wide range of math and science and general reasoning benchmarks, consistent with the types of queries required. The user’s license did not permit Google to use data provided by a user. We also selected a local version of OpenAI’s ChatGPT-4o^15^, another multimodal model with math data included for training. The local version, PittGPT, adds protections that ensure data input to the model will be repurposed, and reduces the cost to Pitt users of the system, among other improvements.

Each protocol was uploaded into a LLM as a PDF and evaluated separately. Each question was refined throughout query development to ensure reliability. LLM output is randomized, so tests were performed three times to determine effectiveness. Initial approaches included developing context by loading entire legal, ethical, and policy documents for IACUC; however, this produced results that varied with small changes in wording by experimenters. Having an expert select only context related to specific queries was found to be necessary to produce accurate and robust results. Structuring the queries with paragraph breaks was also found to be important. After significant experimentation, the described template was developed, and all refined questions were recast using that template.

### Precision and Recall Metric Testing

Precision and recall offer valuable insight into evaluating the performance of LLMs especially for information retrieval or classification tasks. Precision measures the proportion of identified inconsistencies that are true inconsistencies while recall measures the proportion of actual inconsistencies that are correctly identified by the LLM.

Fifty new IACUC protocol samples were collected, and each query ran on all samples, computing precision and recall. Tables for each sample were constructed and LLM outputs for each query were categorized as either true positives, true negatives, false positives, or false negatives. Each protocol was manually reviewed to validate the accuracy of the LLM-generated responses, and the total counts of positive and negative answers were recorded. To assess the consistency and potential variability in LLM outputs, all queries were performed three times. Furthermore, an F1 score was calculated for both LLMs and for each query across all samples. According to established evaluation criteria, an F1 score of 0.7 characterizes the LLM as average, whereas a score of 0.9 indicates a high-performing model^24^. A table was generated detailing total precision and F1 scores for each query. All tables and calculations were developed using Microsoft Excel (Version 2410 Build 16.0.18129.20200).

## Supporting information

Supplementary Material for Table 2: Detailing Prompt List Structure

## End Notes

### Author Contributions

Brent Vasquez is responsible for designing experiments, formulating prompts, conducting precision and recall metrics, and developing tables and figures. He developed initial manuscript drafts and made significant contributions to writing.

Jeremy DeRicco is responsible for providing IACUC protocols and expertise on the IACUC review process. He helped design experiments, analyze data, and made significant contributions to writing.

Bill J Yates is responsible for expertise on the IACUC review process and helped to design experiments and evaluations. He helped analyze data and made significant contributions to writing. Robert K Cunningham is responsible for providing artificial intelligence and software engineering expertise. He designed the evaluation, helped analyze the data, and made significant contributions to writing.

Supplementary Information is available for this paper.

