## Supplementary Material for Table 2: Detailing Prompt List Structure for "AI for IACUC: Accurate Initial Assessment of Institutional Animal Care and Use Committee Protocols"

| Index | Prompt | Details | **Source:** American Veterinary Medical Association (AVMA), National Institute of Health (NIH) |
|---|---|---|---|
| **Output customization, persona** | Introduction Prompt | Summarize the document. Write all responses to the prompts in a polite and professional tone. Provide initial timestamp for beginning of analysis. | N/A |
| **Missing Information** | 1 | Verify that the Principal Investigator listed in section 3.0 is also listed in either section 7.0 or section 8.0. If this is correct put a green check mark. If it is not correct put a red 'x.' | NIH Guidelines |
| | 5 | Our policy requires that anyone listed as a "DEA Registrant" for a controlled drug must be listed as "General Staff" or "Surgical Staff" under the "Protocol Identification" section. Verify if there is anyone listed as a "DEA Registrant". They must be also found under "General Staff" or "Surgical Staff". If correct put a green check mark. If incorrect put a red 'x'. | NIH Guidelines |
| **Impermissible Information** | 4 | Verify that 'Cost', money, or anything related to monetary value is not mentioned in the response to any of the questions. If this is correct put a green check mark. If this is not correct put a red 'x'. | NIH Guidelines |
| **Mismatched Sections** | 2 | Verify that any staff listed under "Breeding Section 1" is also listed under "general staff" or "surgical staff" on the first page. If this is correct put a green check mark. If this is not correct put a red 'x'. | NIH Guidelines |
| | 3 | Public Health Service (PHS) funding sources include: Administration for Children and Families (ACF); Administration on Aging (AoA); Agency for Healthcare Research and Quality (AHRQ); Agency for Toxic Substances and Disease Registry (ATSDR); Centers for Disease Control and Prevention (CDC); Centers for Medicare & Medicaid Services (CMS); Federal Occupational Health (FOH); Food and Drug Administration (FDA); Health Resources and Services Administration (HRSA); Children's Hospital (CHP foundation); Indian health Service (HIS); National Institutes of Health (NIH); Substance Abuse and Mental Health Services Administration (SAMHSA); and the Public Health Service Commissioned Corps. If the only funding source checked in the document is a PHS funding source, then there should be a 'yes' response under "Is this study funded in part or whole by a PHS Agency?". If both a PHS funding source and "Internal" are checked then the response under "Is this study funded in part or whole by a PHS Agency? should be 'yes'. If the funding source mentioned in the funding source title table under section 1.3 is not a PHS agency then there should be a 'no' response under "Is this study funded in part or whole by a PHS agency?". If this is correct put a green check mark, if this is incorrect put a red 'x'. | NIH Guidelines |
| | 7 | What is the "number" in the Animal Species Strain and Numbers section 1.0 table? Verify that the numbers listed in the first three fields, 'Weaned & Adult Animals Used Experimentally', 'Suckling Animals Used Experimentally', and 'Breeders Not Used Experimentally' in the table in section 3.0 are not greater than the "number" in the Animal Species Strain and Numbers section 1.0 table. Do not include 'Animals euthanized prior to weaning & not used experimentally'. Show me the calculation. If this is correct put a green check mark. If this is not correct put a red 'x'. | NIH Guidelines |
| **Identification Of Cases That Fail To Follow Standards** | 6 | Chemical agents and drugs typically used in animal research include anesthetic agents (like isoflurane, ketamine), toxins (like ricin, botulinum toxin), carcinogens (like DMBA), mutagens (like urethane), irritants (used in skin irritation tests), pharmaceuticals (approved by the United States Pharmacopeia and National Formulary (USP-NF)), heavy metals (like lead, mercury), radioactive isotopes, antigens, adjuvants, immunosuppressants, neurotoxins, specific chemicals like hormones, neurotransmitters, opioids, adeno-associated virus (AAV) vector, peptides, or metabolic substrates may also be used. Any agents or drugs not found from this list are incorrect. In the "chemical agents exposure" and "drug" sections what are all the chemical agents and drugs? What agents or drugs do not match the list? Put a green check mark next to each one that is correct. Put a red 'x' next to each one that is incorrect. | NIH Guidelines |
| | 8 | Our policy follows the IACUC guidelines for USDA pain and distress classifications. In the 'Pain and Distress Classification' section, identify which category (Cat B, Cat C, Cat D, Cat E) has a checkmark directly underneath it and make sure to confirm which category has a checkmark closest to it. Count the total number of checkmarks. Identify the species associated with the categories. Verify if the categories with checkmarks directly underneath them match the description provided in section 2.0. If this is correct put a green check mark. If incorrect put a red 'x'. | AVMA and NIH Guidelines |
| | 9 | Our policy for rats requires that if the age genotyped is less than or equal to 14 days then no agent is needed and should never be mentioned, but if age genotyped is greater than 16 days then general anesthesia is administered and if the age genotyped is greater than 28 days then anesthesia and analgesia is administered. Our policy for tail clipping mice requires that if the age genotyped is less than or equal to 16 days then no agent is needed and should never be mentioned, but if age genotyped is greater than 16 days then anesthesia is administered and if the age genotyped is greater than 28 days then anesthesia and analgesia is administered. This policy only applies to mice and rat. Any other species is incorrect. Anesthesia and analgesia should be correct according to the national institute of health (NIH). | AVMA |
| | 10 | In "Breeding Section 2 (genotyping/euthanasia/numbers)" section 1.0 what is the "species" and "age genotyped"? Is the type of anesthesia or analgesia appropriate, and correct? If correct show reasoning and end with a green check mark. If not correct put a red 'x'. | AVMA |
| | 11 | Our policy follows the methods of euthanasia according to the American Veterinary Medical Association (AVMA) guidelines. Are the methods of euthanasia under the Final disposition section 1.0 in the protocol justified according to our policy? If this is correct put a green check mark. If this is not correct put a red 'x'. | AVMA |
| **Output customization, persona** | Concluding Prompt | List all prompts with a red 'x' and add them up. Then list all prompts with a green check mark and add them up. Provide final timestamp for end of analysis. | N/A |
